# Mechanistic Identifiability Preservation for Hybrid Neural Differential Equations

**DOI:** 10.1101/2024.12.08.627408

**Authors:** Benjamin Whipple, Esteban A. Hernandez-Vargas

## Abstract

Hybrid neural differential equations (HNDEs) embed neural network components within mechanistic scaffolds, combining the structural interpretability of domain-derived models with the approximation power of neural dynamics. Despite their growing adoption in biology and engineering, neural augmentation can introduce observational degeneracies that compromise mechanistic identifiability and scientific interpretability.

In this paper, we develop a theoretical framework for practical preservation of mechanistic identifiability in HNDEs. We formalize bounded neural correction classes and derive Gronwall-type trajectory and observational discrepancy bounds linking neural perturbations to mechanistic parameter ambiguity. We further establish sufficient conditions under which hybrid neural corrections preserve approximate mechanistic parameter recoverability up to explicitly quantifiable tolerances.

Empirical likelihood profile analyses on benchmark systems confirm that neural augmentation systematically weakens—but does not eliminate—mechanistic identifiability, revealing a fundamental expressiveness–identifiability trade-off. These results provide theoretical foundations and actionable guidance for deploying HNDEs in scientific intelligent computing.

## 1. INTRODUCTION

Neural dynamical systems—systems of differential equations whose right-hand sides are parameterized by neural networks—have become fundamental tools in intelligent computing [1, 2]. The conceptual generalization to learned dynamics—where the neural vector field is fitted from data rather than prescribed—underlies modern NDEs [3]. This perspective has been extended to scientific computing, where the task is the discovery and simulation of complex dynamical phenomena from incomplete data [4, 5].

Scientific machine learning faces a distinctive challenge: models must be both predictively accurate and mechanistically interpretable [6, 7]. In biology and engineering, purely data-driven approaches lack the structural constraints needed for scientific inference, while purely mechanistic models become intractable when system interactions are only partially characterized [8, 9]. Hybrid neural differential equations (HNDEs) address this tension by embedding neural network components within a mechanistic differential equation models [4, 10, 11].

HNDEs have antecedents in the grey-box process models of the 1990s, where neural networks were used to correct incompletely specified mechanistic descriptions [12, 13, 14, 15]. Modern formulations integrate the neural component directly into the ODE right-hand side rather than as an additive error term [4, 16, 17]. Applications span ecological parameter recovery [18, 17], chemical bioprocess engineering [19, 10, 20, 21, 22, 23], and epidemiological compartment discovery [24, 25, 26, 27]. The term “universal differential equation” of [4] emphasizes the universality of the neural approximator; we prefer HNDE following [12] to emphasize the hybrid mechanistic and data-driven character.

Parameter identifiability is a classical concern in mechanistic modeling [28, 29, 30]. Structural identifiability analysis determines whether parameters are theoretically recoverable from noise-free observations, while practical identifiability methods—Fisher information matrices and likelihood profiles—assess recoverability from finite, noisy data [31]. The inclusion of flexible neural components introduces new degeneracies that are not captured by classical structural analysis.

Two critical challenges limit the scientific utility of HNDEs. First, calibration is sensitive to hyper-parameter choices and principled guidance on their relative importance is lacking. Second, incorporating a flexible neural component may compromise the identifiability of mechanistic parameters, which could otherwise lead to overfitting and loss of interpretability [32]. Previous work [11] enforced identifiability during training via a meta-optimization procedure that treats mechanistic parameters as hyper-parameters and applies local Hessian-based identifiability diagnostics. Uncertainty estimates have been explored for the mechanistic and neural components [33].

This paper complements previous contributions on hybrid neural differential equations [11, 32, 33] by developing a theoretical framework that characterizes how neural augmentation affects mechanistic identifiability and explicitly links neural network architecture to identifiability preservation. In contrast to primarily workflow-oriented or empirical studies, we derive sufficient conditions for bounded identifiability degradation and employ likelihood profiles to evaluate mechanistic identifiability [31]. Figure 1 illustrates the principal integration modes considered in hybrid neural dynamical systems. In this work, we focus specifically on Cases I and II: learning unknown kinetic mechanisms and approximating unmeasured intermediate states.

**Figure 1:**
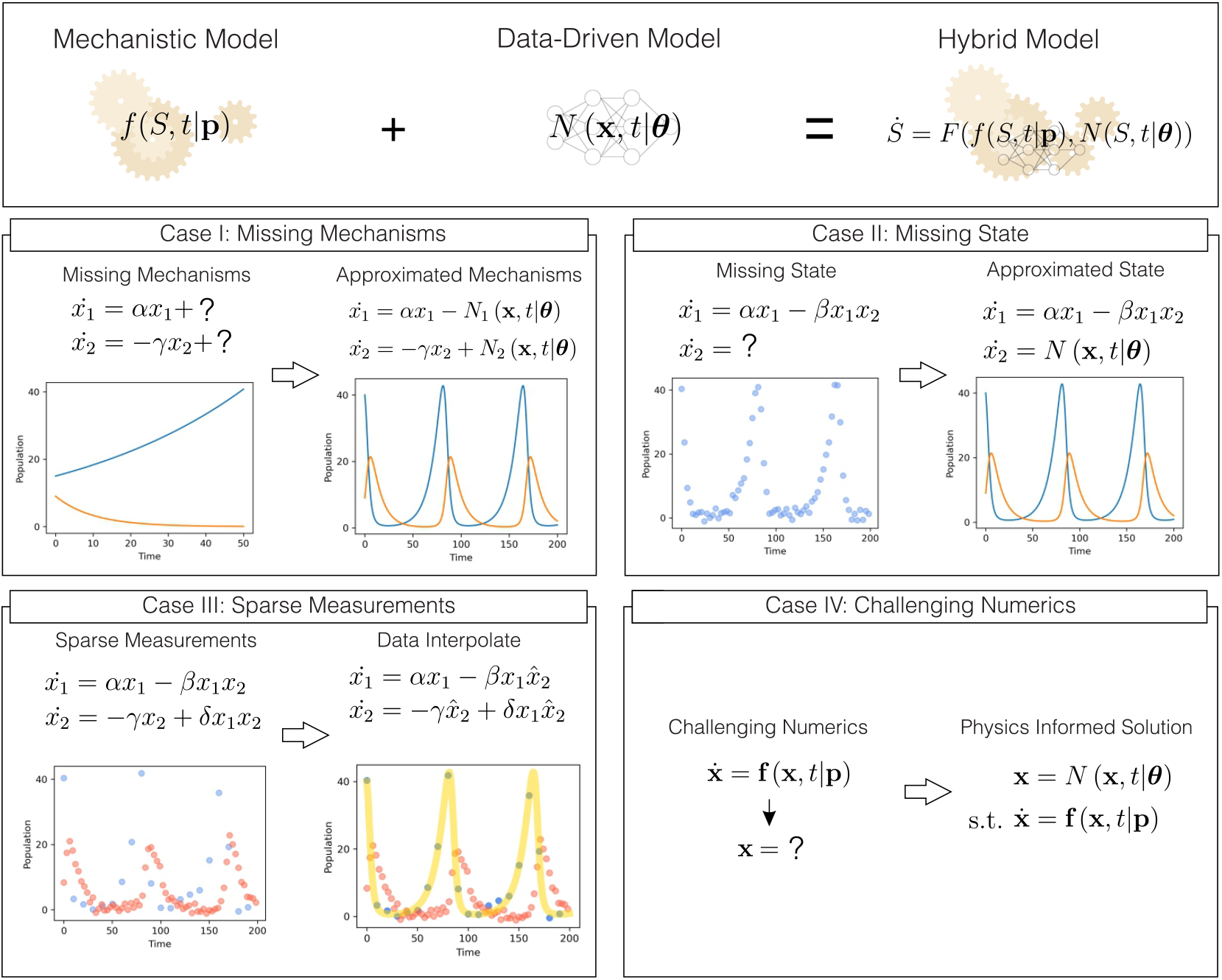
Principal modes of neural–mechanistic integration illustrated with the Lotka– Volterra model. Case I: neural component *N* approximates unknown interaction dynamics. Case II: *N* approximates an unmeasured but mechanistically necessary state. Case III: machine learning interpolates low-frequency measurements to improve calibration. Case IV: neural methods address challenging numerical problems such as PDEs.

To sum up, this paper makes the following four contributions:

i. Hybrid neural differential equations are extended to incorporate theorized but unmeasured intermediate states, enabling neural augmentation under partial mechanistic observability (Section 3).
ii. A systematic hyper-parameter sensitivity analysis is performed, showing that batch horizon, learning rate, and training iterations are the dominant factors governing calibration performance, while network width and batch size have comparatively limited influence.
iii. A theoretical framework for mechanistic identifiability in universal differential equations is developed. In particular, the notion of preservation of mechanistic identifiability is formalized, a Gronwall-based sufficient condition for bounded identifiability degradation is established (Section 2.1). To our knowledge, this provides one of the first mathematical frameworks characterizing identifiability preservation in hybrid neural differential equations.
iv. Likelihood profile analyses on glycolysis and Lotka–Volterra benchmark systems demonstrate that neural augmentation systematically weakens—but does not eliminate—mechanistic identifiability, revealing an expressiveness– identifiability trade-off in hybrid neural dynamical systems.

The remainder of this paper is organized as follows. Section 2 introduces the hybrid neural differential equation framework and calibration methodology. Section 2.1 develops the theoretical framework for mechanistic identifiability preservation. Section 3 describes the benchmark systems and experimental setup. Section 4 presents the hyper-parameter and identifiability analyses. Section 5 discusses the implications of neural augmentation for mechanistic interpretability and scientific machine learning.

## 2. HYBRID NEURAL DYNAMICAL SYSTEMS

Neural differential equations (NDEs) provide a continuous-time representation of neural dynamics in which the temporal evolution of the hidden state is governed entirely by a neural network:

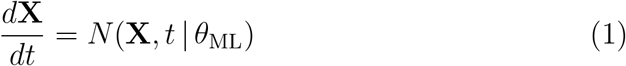

where **X** ∈ ℝ^*n*^ denotes system states, *N*: ℝ^*n*^ × ℝ^+^ × ℝ^*q*^ → ℝ^*n*^ is a neural network parameterized by *θ*_ML_ ∈ ℝ^*q*^, and *t* ∈ ℝ^+^ is time [3].

Incorporating mechanistic prior knowledge through a structured dynamical component *g*: ℝ^*n*^ × ℝ^+^ × ℝ^*p*^ → ℝ^*m*^ parameterized by the mechanistic parameters *θ*_Mech_ ∈ ℝ^*p*^, yields a hybrid neural differential equation (HNDE), in which mechanistic dynamics are augmented by a neural component:

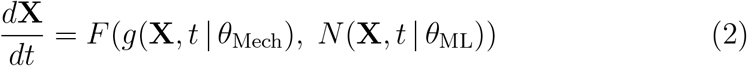

where *F*: ℝ^*m*^ × ℝ^*n*^ → ℝ^*n*^ defines the interaction between the mechanistic and neural components.

Throughout this paper, we focus on the additive HNDE formulation, where the mechanistic and neural components contribute through a super-position structure. In this setting, the general coupling operator *F* in equation (2) is specialized to an additive decomposition,

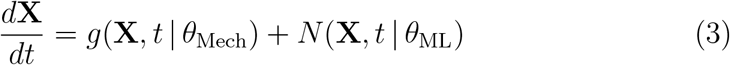

where the mechanistic term *g* captures the known or hypothesized system dynamics, while the neural component *N* represents unresolved, latent, or data-driven effects. This additive structure preserves an explicit separation between mechanistic knowledge and learned corrections, which is advantageous both conceptually and mathematically.

### Remark 1.

*From an analytical perspective, the additive form enables the neural contribution to be interpreted as a structured perturbation of the mechanistic dynamics. This facilitates the study of qualitative properties since the mechanistic backbone remains explicitly accessible. In contrast, more general nonlinear couplings may obscure the mechanistic contribution and complicate the interpretation of parameter effects and dynamical mechanisms*.

### 2.1. Identifiability Preservation Theory

Parameter identifiability—the ability to uniquely recover model parameters from observable data—is a fundamental requirement for scientific interpretability, mechanistic validation, and reliable prediction in dynamical systems [28, 29]. Without identifiability, multiple parameter configurations may generate indistinguishable trajectories, preventing causal interpretation of the underlying biological mechanisms. This challenge becomes particularly critical in HNDEs, where the flexibility of neural components may absorb or mask mechanistic effects, potentially compromising the interpretability of the mechanistic backbone.

For additive HNDEs of the form (3), a central question is therefore whether the introduction of the neural correction term preserves the recoverability of the mechanistic parameters. The additive structure plays an essential role in this context, as it maintains an explicit decomposition between the known mechanistic dynamics and the learned residual component. This separation enables the neural term to act as a constrained correction rather than a complete replacement of the governing dynamics, thereby facilitating the analysis of parameter recoverability.

Following the framework of [32], we adapt the notion of identifiability to hybrid systems by focusing specifically on the mechanistic parameters *θ*_Mech_. In particular, *θ*_Mech_ is said to be identifiable if equality of observable trajectories implies equality of the mechanistic parameters, independently of equivalent neural parameterizations satisfying 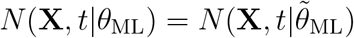.

We do not investigate the identifiability of *θ*_ML_, since neural network parameterizations are well known to admit substantial non-uniqueness due to symmetries, overparameterization, and functional equivalence classes. To formalize this notion, we introduce the concept of preservation of mechanistic identifiability.

#### Definition 1

(Practical Preservation of Mechanistic Identifiability). *Consider system (3) with state* **X**(*t*) ∈ ℝ^*n*^, *mechanistic parameters θ*_Mech_ ∈ Θ_Mech_ ⊆ ℝ^*p*^, *and neural parameters θ*_ML_ ∈ Θ_ML_ ⊆ ℝ^*q*^. *Let*

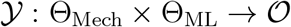

*denote the observation operator mapping parameter pairs to observable trajectories, and let* ∥ · ∥_𝒪_ *denote a norm on the observation space. Assume that the neural component belongs to an admissible class* 𝒩_*M*_ *satisfying*

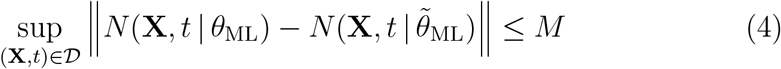

*for some bounded domain* 𝒟⊆ ℝ^*n*^ ×ℝ^+^ *and constant M >* 0. *The system (3)* practically preserves mechanistic identifiability *if, for every ε >* 0, *there exist δ >* 0 *and M >* 0 *such that, for all*

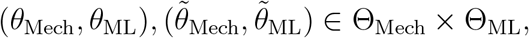

*satisfying the above bounded neural discrepancy condition*,

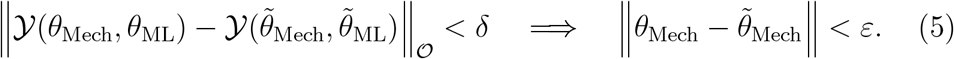

Definition 1 extends the classical notion of practical identifiability to hybrid neural differential equations through an *ε*–*δ* characterization of mechanistic parameter recovery under bounded neural discrepancy. This relaxation is particularly important in HNDEs, since the machine learning component inherently introduces approximation uncertainty through latent dynamics, unresolved interactions, model mismatch, and data-driven corrections. Consequently, exact recovery of the mechanistic parameters is generally unrealistic, even when the mechanistic structure is informative.

#### Remark 2.

*The parameter δ quantifies the observational resolution under which two trajectories are considered practically indistinguishable, while ε measures the induced uncertainty in the associated mechanistic parameterization. The bounded neural discrepancy condition in (4) plays a central role by constraining the admissible variability of the neural correction term over the domain* 𝒟. *Rather than enforcing equality of neural parameterizations, which is generally unrealistic due to the intrinsic non-identifiability and over-parameterization of neural networks, the definition controls the functional discrepancy between neural components*.

Under this framework, the neural component is permitted to absorb unresolved or latent dynamics while remaining sufficiently constrained to avoid large observational degeneracies or fundamentally different mechanistic explanations. Therefore, an additive HNDE framework practically preserves mechanistic identifiability when uncertainty introduced by the neural correction term propagates in a controlled manner to the mechanistic parameters, maintaining the interpretability and scientific relevance of the mechanistic backbone despite the flexibility of the machine learning component.

Before stating the main identifiability theorem, we establish a constructive bound showing how bounded discrepancies between neural correction terms propagate to the observable trajectories of the additive HNDE (3).

#### Proposition 1

(Gronwall-Type Bound for Neural Discrepancies). *Suppose that g*(·, *t* | *θ*_Mech_) *is Lipschitz continuous with constant L*_*g*_ *>* 0, *uniformly in t and θ*_Mech_, *along the trajectories considered:*

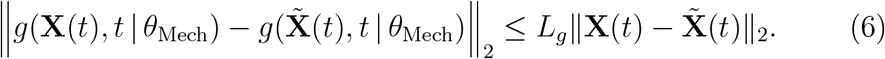

*Assume also that the neural component is Lipschitz continuous in* **X** *with constant L*_*N*_ *>* 0, *uniformly in t and θ*_ML_:

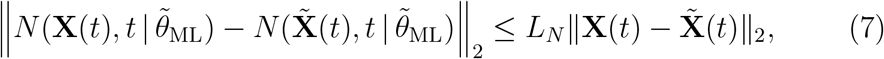

*and assume that the neural discrepancy satisfies* (4) *Let* **X**(*t*) *and* 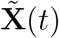 *denote solutions of system (3) associated with* (*θ*_Mech_, *θ*_ML_) *and* 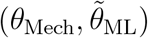, *respectively, with the same initial condition. Then, for every finite horizon T >* 0 *such that* 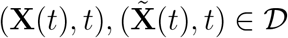*for all t* ∈ [0, *T*],

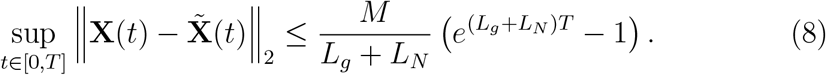

*Assume further that observations are generated by a linear map*

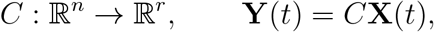

*and collected at times* 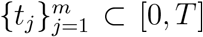. *If* 𝒪 *is equipped with the* 𝓁^*∞*^*-norm over observation times, then*

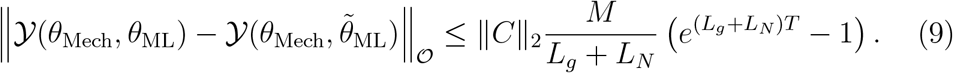

*Proof*. Let

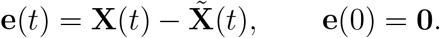

Subtracting the two additive HNDEs in equation (3) gives

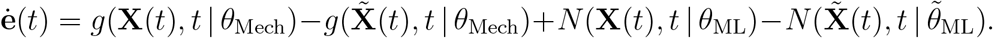

Add and subtract 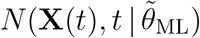to obtain

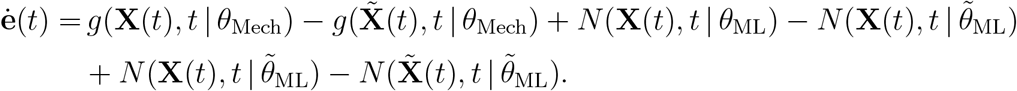

Using equations (6), (7), and (4), we obtain

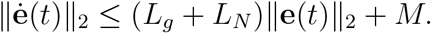

Since **e**(0) = **0**, we have 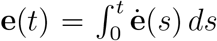*ds*. Taking norms and applying the triangle inequality yields

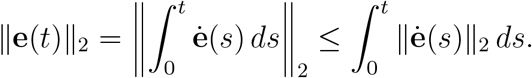

Using ∥**ė** (*s*)∥_2_ ≤ (*L*_*g*_ + *L*_*N*_)∥**e**(*s*)∥_2_ + *M*, we obtain

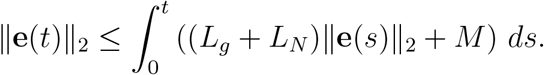

Applying Gronwall’s inequality gives

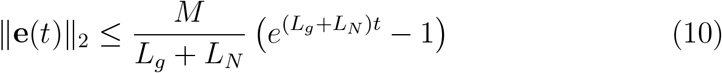

Taking the supremum over *t* ∈ [0, *T*] yields equation (8). The observation bound follows from

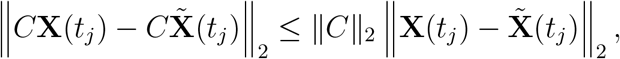

taking the maximum over *j* = 1, …, *m*, and applying equation (8).

#### Remark 3.

*Proposition 1 establishes that bounded discrepancies between neural correction terms induce controlled perturbations in both the hidden trajectories and the associated observable outputs of the additive HNDE (3). The bound* (10) *reveals three fundamental factors governing practical preservation of mechanistic identifiability: the magnitude of the neural discrepancy M, the combined dynamical sensitivity encoded by the Lipschitz constants L*_*g*_ *and L*_*N*_, *and the observation horizon T. Larger neural discrepancies or highly sensitive dynamics amplify trajectory uncertainty, while shorter observation windows and smaller neural corrections improve stability of the mechanistic interpretation*.

Proposition 1 motivates the following theorem, which connects bounded observational perturbations with practical preservation of mechanistic identifiability. More specifically, the theorem shows that if the underlying mechanistic model satisfies a quantitative separation condition, then bounded neural corrections cannot induce arbitrarily large ambiguities in the mechanistic parameterization.

#### Theorem 1

(Bounded Neural Correction and Practical Identifiability Preservation). *Consider the additive HNDE (3) with hybrid observation operator 𝒴 and purely mechanistic observation operator𝒴_0_. Assume:*

i. ***Mechanistic separation***. *There exists c >* 0 *such that*

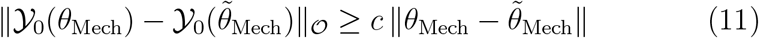

*for all* 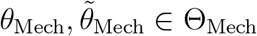 _Mech_.
ii. ***Bounded hybrid deviation***. *There exists η* ≥ 0 *such that*

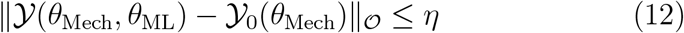

*for all* (*θ*_Mech_, *θ*_ML_) ∈ Θ_Mech_ × Θ_ML_.

*If two hybrid observations satisfy*

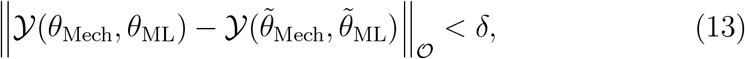

*then*

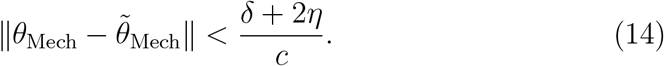

*In particular, if δ* = 0 *and η* = 0, *then* 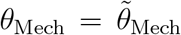, *and the hybrid system exactly preserves mechanistic identifiability*.

*Proof*. By the mechanistic separation condition,

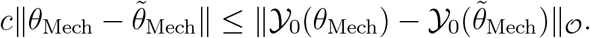

Adding and subtracting the hybrid observations gives

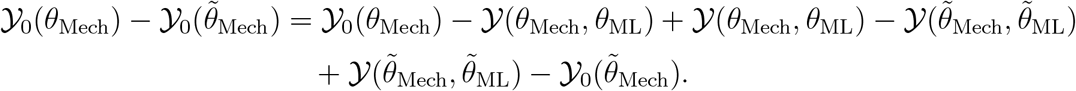

Taking norms and applying the triangle inequality yields

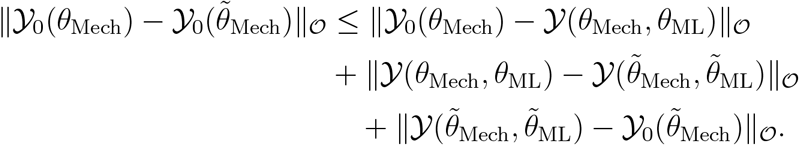

Using (12) on the first and third terms and (13) on the middle term gives

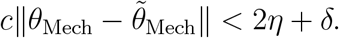

Therefore,

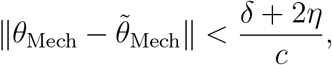

which proves (14).

#### Remark 4.

*The bounded hybrid deviation condition (12) is the key assumption underlying approximate preservation of mechanistic identifiability. It guarantees that the neural correction term acts as a controlled perturbation of the purely mechanistic observation operator rather than an unrestricted replacement of the mechanistic dynamics. Consequently, the hybrid model remains sufficiently close to the mechanistic representation for parameter distinguishability to persist up to an explicitly quantifiable tolerance*.

## 3. EXPERIMENTAL SETUP

We evaluate on two benchmark biological systems that represent complementary sources of model uncertainty: a missing kinetic mechanism (glycolysis) and a missing state measurement (three-species Lotka–Volterra). Both systems are simulated to generate synthetic datasets with additive multiplicative Gaussian noise (*µ* = 1, *σ* = 0.05).

### 3.1. Model calibration

Both NDE and HNDE models are calibrated using gradient-based methods, with gradients computed via automatic differentiation through a compatible ODE solver [34, 35, 36]. We employ the ADAM optimizer [37], consistent with recent HNDE literature [10, 38].

We adopt the mini-batch loss formulation: short time windows are sampled as initial conditions, integrated forward, and prediction errors accumulated across segments before updating parameters. Mini-batch training is known to improve convergence for time-series neural dynamical models [39, 40, 41] and is used throughout with mean absolute error (MAE).

### 3.2. Benchmark 1: Missing mechanism (glycolysis)

Glycolysis in yeast is a widely used benchmark for system identification [42, 10, 43, 44]. Following [10], we treat the *S*_1_ → *S*_2_ kinetic term as unknown:

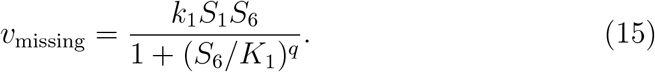

Setting *v*_missing_ = 0 renders the incomplete system incapable of reproducing the dynamics [10]. The full system and NDE formulations are shown in Figure 2. Simulation parameters: *J*_0_ = 2.5, *k*_1_ = 100, *k*_2_ = 6, *k*_3_ = 16, *k*_4_ = 100, *k*_5_ = 1.28, *k*_6_ = 12, *k* = 1.8, *κ* = 13, *q* = 4, *K*_1_ = 0.52, *ψ* = 0.1, *N* = 1, *A* = 4; initial conditions (*S*_1_, …, *S*_7_)(0) = (1.6, 1.5, 0.2, 0.35, 0.3, 2.67, 0.1); time horizon *t* ∈ [0, 6]; 225 observations [10]. The initial guess for the unknown HNDE is ***θ***_Mech_ = [1, 1, 10, 100, 1, 10, 1, 10, 0.1, 1, 1]. The first 150 observations form the training set; the remaining 75 the test set.

**Figure 2:**
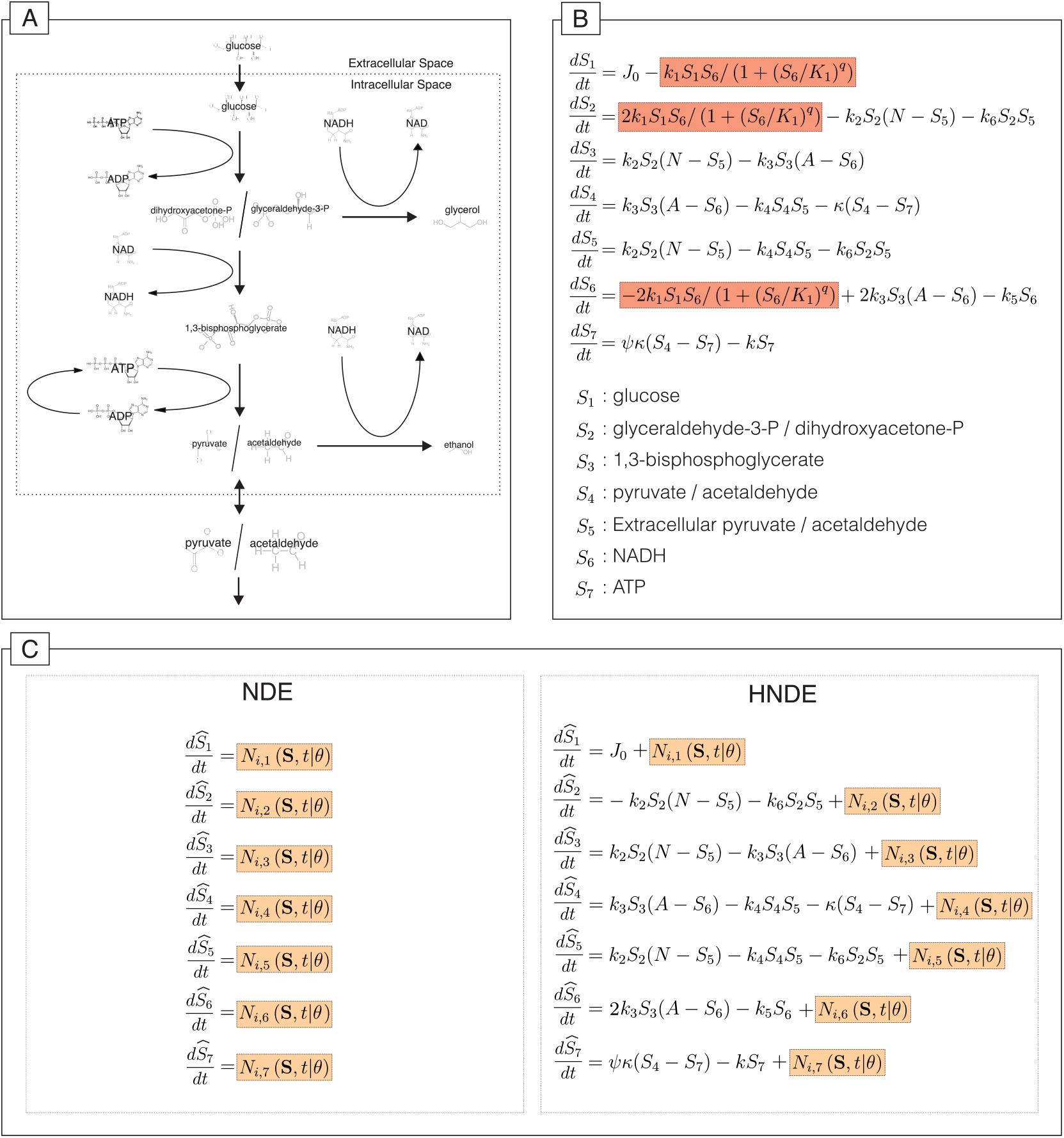
NDEs applied to missing mechanistic knowledge in the yeast glycolysis model [44].(A) Reaction network. (B) Model equations with missing term highlighted. (C) NDE and HNDE formulations; neural entry points highlighted.

### 3.3. Benchmark 2: Missing state (Lotka–Volterra)

We consider a three-species Lotka–Volterra model in which the primary prey *S*_1_ is unmeasured but mechanistically necessary for the dynamics of the intermediate species *S*_2_. System equations and NDE formulations are shown in Figure 3.

**Figure 3:**
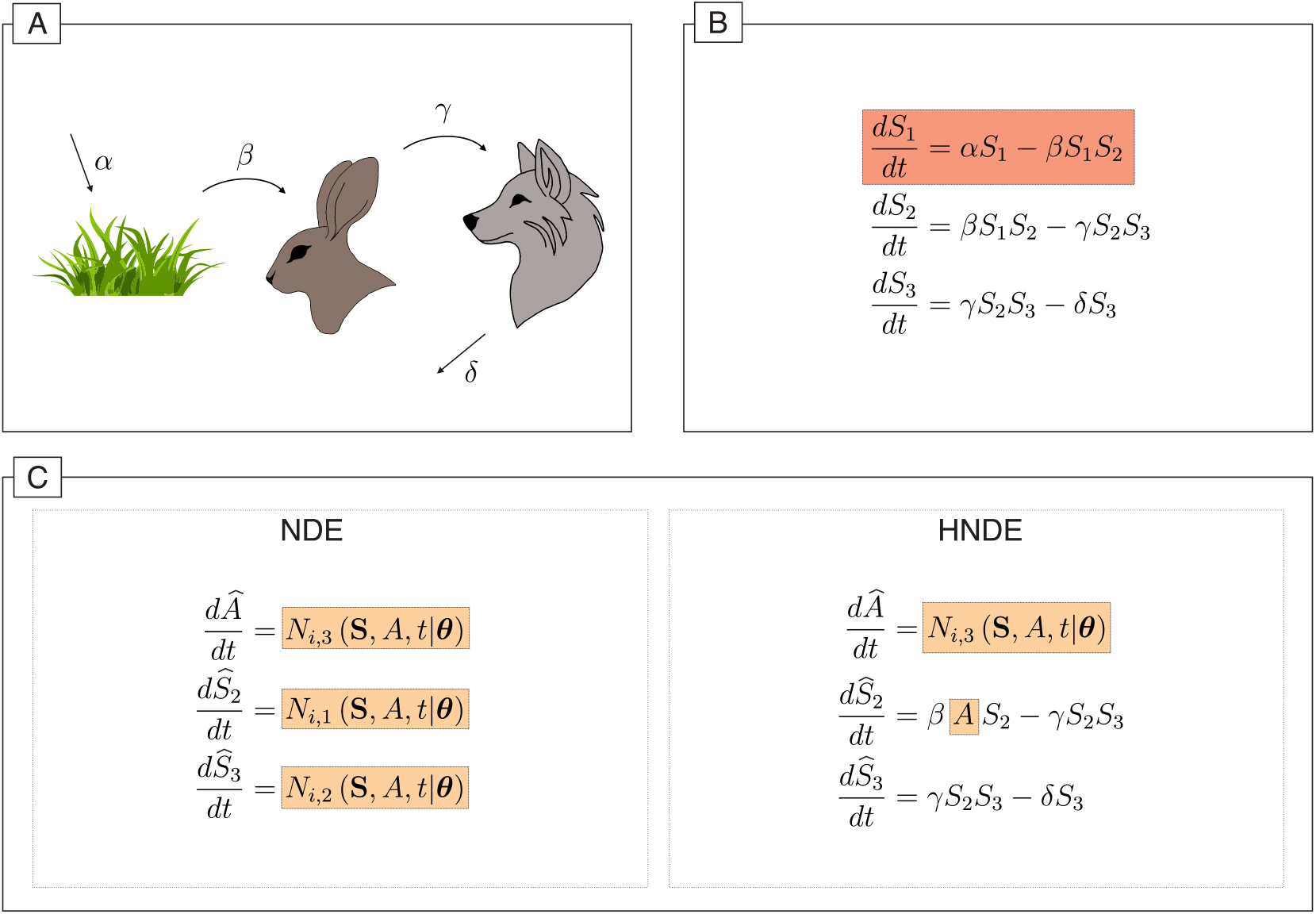
NDEs applied to missing state knowledge in a three-species Lotka–Volterra system. (A) Interaction diagram. (B) Model equations with missing state highlighted.(C) NDE and HNDE formulations; neural entry points highlighted.

Simulation parameters: *α* = 0.5, *β* = 1.5, *γ* = 3.0, *δ* = 1.0; initial conditions *S*_1_(0) = *S*_2_(0) = *S*_3_(0) = 1.0; time horizon *t* ∈ [0, 15]; 150 observations. The initial guess for the unknown HNDE is ***θ***_Mech_ = [1.0, 2.0, 1.0]. The first 90 observations serve as training data; the final 60 as test data.

### 3.4. Hyper-parameter study design

We conduct a factorial experiment over the levels in Table 1. Nine replicates are generated per combination, calibrated using ADAM. Results are analyzed with the fixed effects model

**Table 1:**
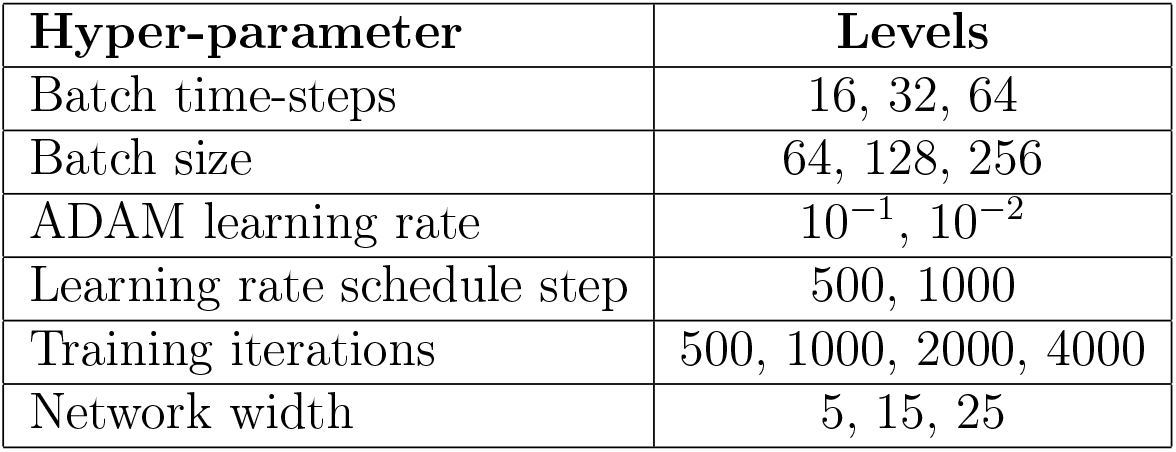
Hyper-parameter levels used in the factorial experiment.

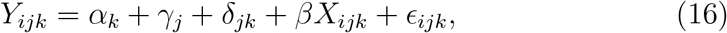

implemented in R [45], with training accuracy as the response to prevent information leakage [38]. Hyper-parameters are coded as categorical variables with polynomial contrasts; fixed effects control for test problem (*α*), model structure (*γ*), and their interaction (*δ*). ANOVA decomposes variance attributable to each hyper-parameter and first-order interactions. All three model structures (pure NDE, known HNDE, unknown HNDE) are included simultaneously for both benchmark problems.

### 3.5. Identifiability analysis design

We assess practical identifiability via likelihood profiles for both benchmark systems [31]. Each mechanistic parameter *θ*_*i*_ is held fixed at 11 values spanning 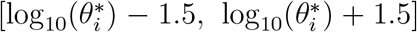 while the remaining parameters are re-optimized; 20 replicates per value account for stochastic calibration. Quadratic smoothing splines summarize profiles across replicates. Fully mechanistic reference profiles—3 replicates, differential evolution on the full dataset—provide an upper bound on achievable identifiability. All analyses use a 5-neuron width with optimal training hyper-parameters determined from the factorial study.

### 3.6. Implementation

All models were implemented in Python using PyTorch and TorchDiffeq [46, 47]. Neural networks consist of a single hidden layer with `tanh` activation, selected via preliminary experiments for sufficient expressive power, numerical stability, and training efficiency. For unknown HNDE models, *θ*_Mech_ is estimated jointly with *θ*_ML_ via ADAM with multiplicative learning rate scheduling. L-BFGS polishing was found to yield negligible improvement at meaningful computational cost and was excluded. Each model structure was evaluated across network widths {5, 15, 25} with 50 replicates per configuration for the predictive performance study. All experiments were run on an Intel i5-12600KF CPU.

## 4. NUMERICAL RESULTS

Batch horizon, learning rate, and training iterations are the dominant determinants of HNDE calibration performance (Table 2). Model structure and its interaction with the test problem also explain substantial variance, reflecting the problem-specific suitability of each approach. In contrast, network width, batch size, and learning rate schedule have negligible effects.

**Table 2:** ANOVA results for the hyper-parameter factorial experiment (fixed effects model, equation 16). All NDE and HNDE structures across both benchmark problems are included. The five most influential variables are bolded.

| Variable | Df | Sum Sq | Mean Sq | F value | Pr(>F) |
| --- | --- | --- | --- | --- | --- |
| Problem | 1 | 0.18 | 0.18 | 1.97 | 0.1600 |
| **Structure** | 2 | 585.85 | 292.92 | 3277.88 | 0.0000 |
| Size | 2 | 4.13 | 2.07 | 23.13 | 0.0000 |
| **Batch Time** | 2 | 23.83 | 11.92 | 133.35 | 0.0000 |
| Batch Size | 2 | 0.73 | 0.37 | 4.11 | 0.0165 |
| **Learning Rate** | 1 | 166.52 | 166.52 | 1863.38 | 0.0000 |
| Learning Rate Step | 1 | 5.69 | 5.69 | 63.63 | 0.0000 |
| **Iterations** | 3 | 69.29 | 23.10 | 258.47 | 0.0000 |
| **Problem**×**Structure** | 2 | 691.90 | 345.95 | 3871.30 | 0.0000 |
| Size×Batch Time | 4 | 11.90 | 2.98 | 33.30 | 0.0000 |
| Size×Batch Size | 4 | 0.24 | 0.06 | 0.69 | 0.6021 |
| Size×Learning Rate | 2 | 11.80 | 5.90 | 66.00 | 0.0000 |
| Size×Learning Rate Step | 2 | 2.09 | 1.05 | 11.70 | 0.0000 |
| Size×Iterations | 6 | 1.20 | 0.20 | 2.24 | 0.0365 |
| Batch Time×Batch Size | 4 | 0.21 | 0.05 | 0.59 | 0.6710 |
| Batch Time×Learning Rate | 2 | 2.02 | 1.01 | 11.32 | 0.0000 |
| Batch Time×LR Step | 2 | 1.80 | 0.90 | 10.06 | 0.0000 |
| Batch Time×Iterations | 6 | 2.83 | 0.47 | 5.27 | 0.0000 |
| Batch Size×Learning Rate | 2 | 1.01 | 0.50 | 5.64 | 0.0036 |
| Batch Size×LR Step | 2 | 0.00 | 0.00 | 0.02 | 0.9847 |
| Batch Size×Iterations | 6 | 0.40 | 0.07 | 0.75 | 0.6059 |
| Learning Rate×LR Step | 1 | 8.91 | 8.91 | 99.72 | 0.0000 |
| Learning Rate×Iterations | 3 | 6.92 | 2.31 | 25.81 | 0.0000 |
| LR Step×Iterations | 3 | 8.10 | 2.70 | 30.22 | 0.0000 |
| Residuals | 18190 | 1625.52 | 0.09 |  |  |

Network width can be fixed to 5–15 neurons, batch size kept low, and learning rate scheduling simplified or omitted without material performance loss. This observation is also consistent with the practical identifiability bound in equation (14), where increasing the flexibility of the neural correction can enlarge the admissible hybrid deviation term *η*, thereby increasing the ambiguity in the recovered mechanistic parameters. Compact neural architectures therefore provide both computational efficiency and improved preservation of mechanistic interpretability.

HNDEs achieve strong predictive performance under both mechanistic and state uncertainty, though the advantage of mechanistic structure depends critically on the nature of the missing knowledge. For the glycolysis system (Figures 4 and 6), all structures achieve broad agreement with test data, with known and unknown HNDE models exhibiting lower trajectory variability. For the three-species Lotka–Volterra system (Figures 5 and 6), the ordering reverses: the pure NDE achieves the best predictive performance, followed by the unknown-parameter HNDE, with the known-parameter HNDE performing worst. This reversal demonstrates that imposed mechanistic structure becomes a constraint rather than an asset when an unmeasured state introduces ambiguity that the fixed structure cannot resolve.

**Figure 4:**
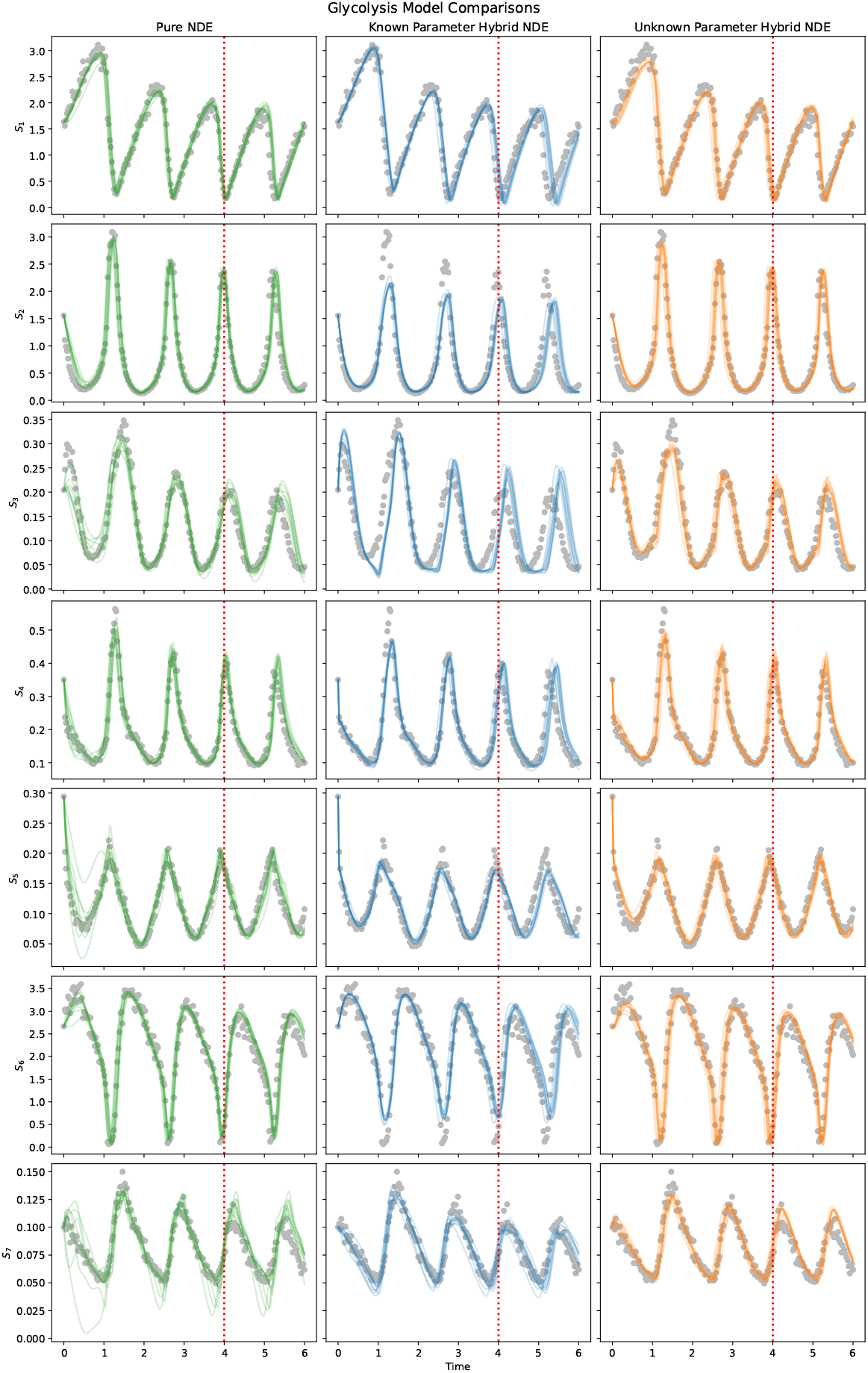
Predictions of 15-neuron models for the glycolysis benchmark. Grey dots: test data; colored lines: 10 best trajectories per structure. Pure NDE predictions exhibit higher variability than HNDE predictions.

**Figure 5:**
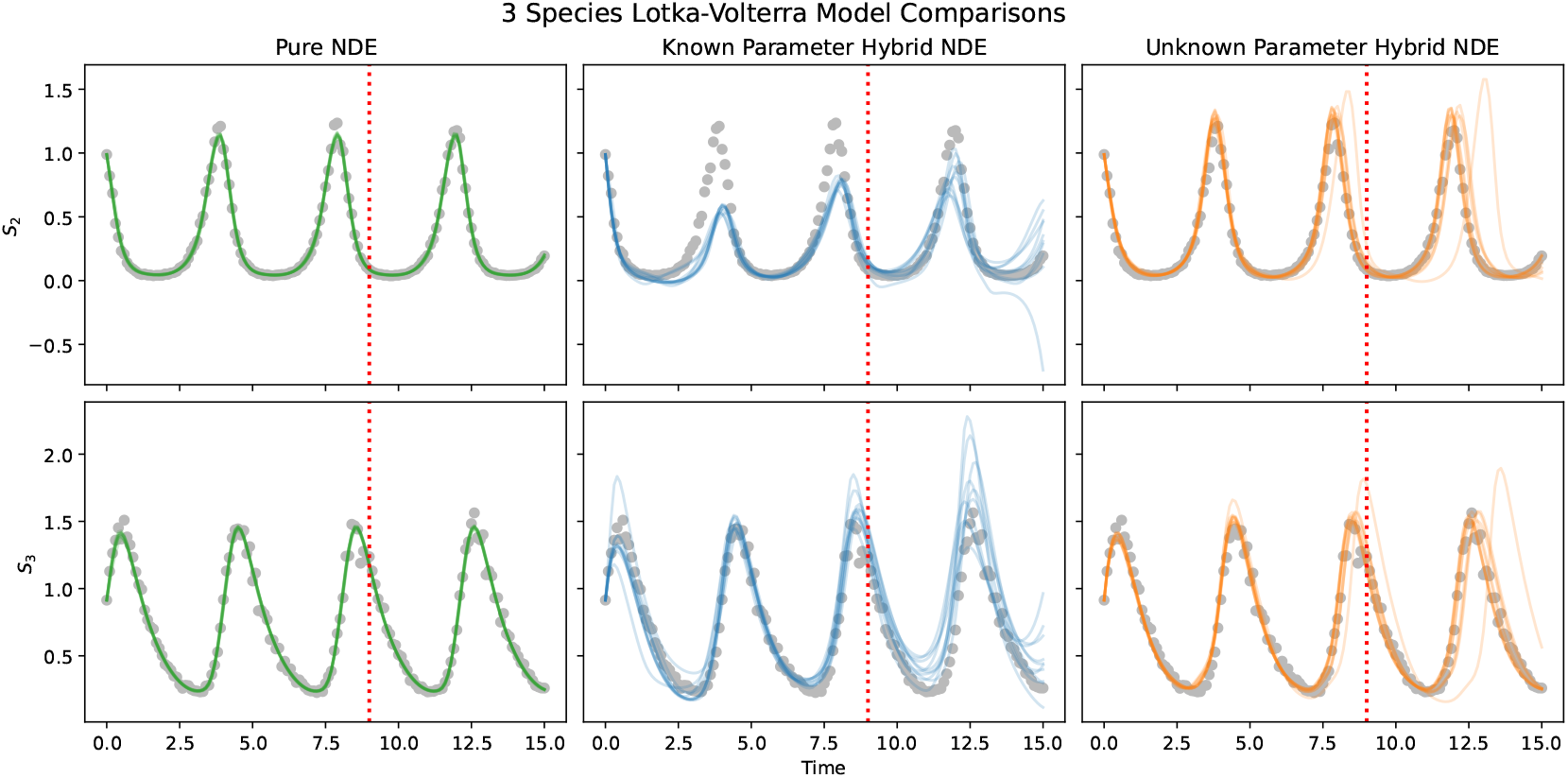
Predictions of 15-neuron models for the Lotka–Volterra benchmark. Grey dots: test data; colored lines: 10 best trajectories per structure. Pure NDE predictions exhibit lower variability than HNDE predictions, reversing the glycolysis pattern.

**Figure 6:**
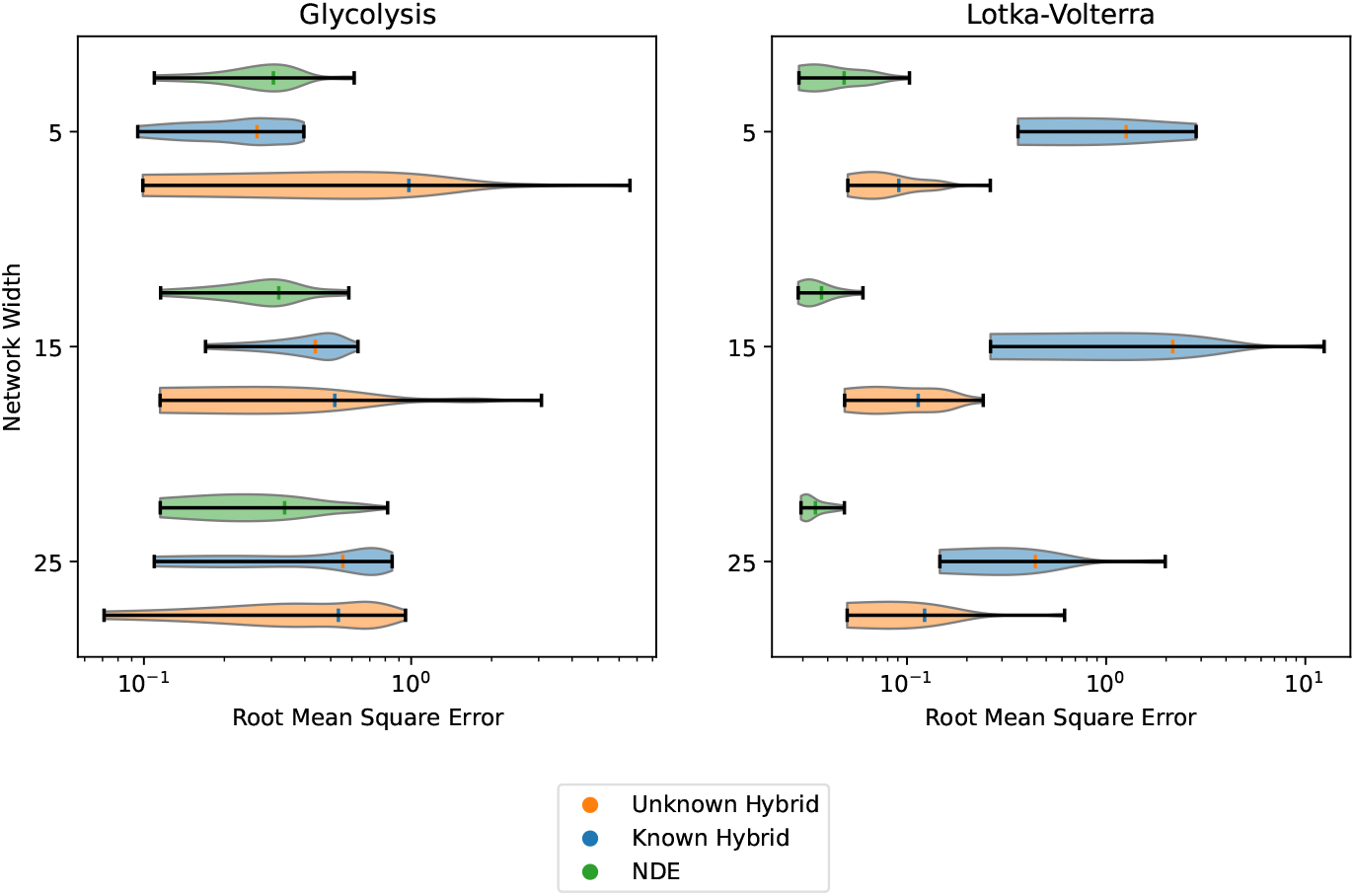
RMSE across replicates for each model structure and network width. All structures perform comparably on glycolysis; on the Lotka–Volterra system, the pure NDE dominates, with HNDE models exhibiting substantially worse performance and higher variability.

Predictive performance is largely invariant to network width across both systems, confirming that 5-neuron networks suffice for these benchmarks [48]. This result further supports the use of analytically tractable compact architectures, since the practical identifiability bound in equation (14) implies that controlling the flexibility of the neural correction term reduces the admissible hybrid deviation *η*, thereby improving preservation of mechanistic parameter recoverability.

Neural augmentation systematically weakens mechanistic parameter identifiability across both benchmark systems, empirically confirming the expressiveness– identifiability trade-off formalized in Definition 1. For the glycolysis model (Figure 7), all parameters are clearly identifiable under the fully mechanistic model. Under the HNDE, only *A* and *k*_6_ retain clear identifiability; *k* and *ψ* become entirely non-identifiable, while remaining parameters show reduced profile convexity. For the Lotka–Volterra model (Figure 8), all parameters are identifiable mechanistically, but the HNDE degrades identifiability of *γ* and *δ* and renders *β* entirely non-identifiable.

**Figure 7:**
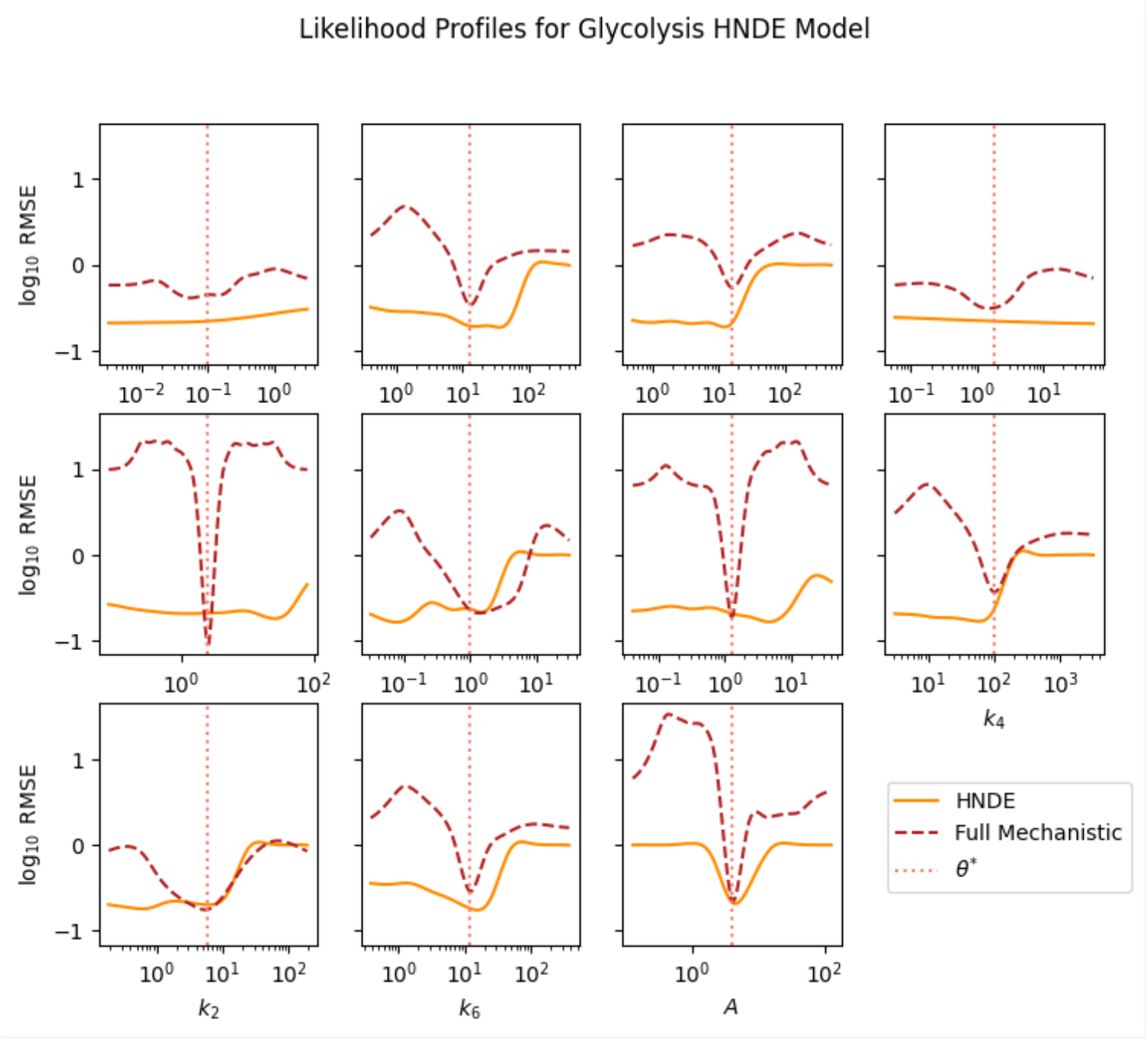
Likelihood profiles for the glycolysis HNDE model. Solid line: HNDE profile; dashed line: fully mechanistic profile; dotted vertical line: true parameter value. Convexity about the true value indicates practical identifiability. All parameters are identifiable under the full mechanistic model; only *k*_6_ and *A* remain clearly identifiable under the HNDE.

**Figure 8:**
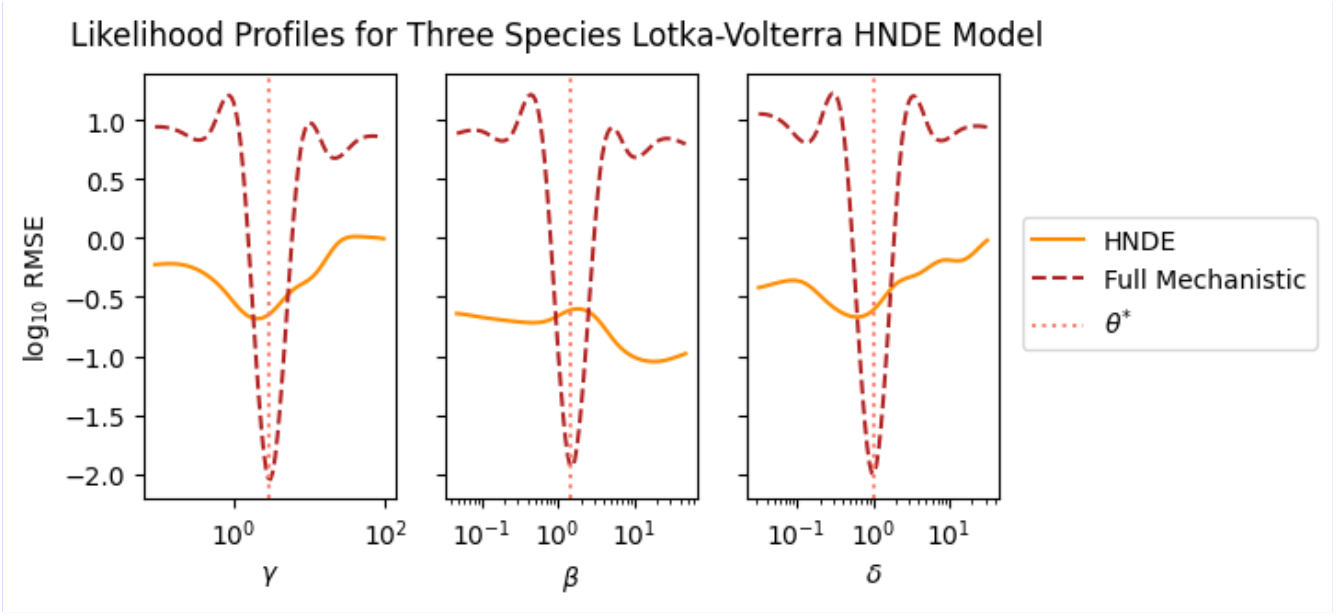
Likelihood profiles for the three-species Lotka–Volterra HNDE model. Solid line: HNDE profile; dashed line: fully mechanistic profile; dotted vertical line: true parameter value. All parameters are identifiable mechanistically; only *γ* and *δ* remain identifiable under the HNDE. The non-identifiability of *β* under the HNDE contrasts sharply with its identifiability under the fully mechanistic model.

Identifiability degradation is partial rather than total: parameters tightly coupled to the retained mechanistic scaffold remain recoverable. This observation is consistent with Theorem 1, where the bounded hybrid deviation term *η* imposes an explicit upper bound on mechanistic parameter ambiguity through equation (14). Consequently, bounded neural corrections may weaken parameter recoverability without fully destroying mechanistic identifiability. These results further validate likelihood profiles as an appropriate diagnostic for HNDE identifiability, confirming their robustness advantage over Fisher Information Matrix methods under stochastic calibration [31].

## 5. DISCUSSION

This paper advanced the theory and practice of hybrid neural dynamical systems for scientific intelligent computing through four contributions. We extended HNDEs to accommodate unmeasured intermediate states, identified the three critical hyper-parameters governing calibration performance, introduced and formalized preservation of mechanistic identifiability with a constructive Gronwall-based sufficient condition. Empirical likelihood profile analyses on glycolysis and Lotka–Volterra benchmarks confirmed that neural augmentation weakens—but does not eliminate—mechanistic identifiability, revealing a fundamental expressiveness–identifiability trade-off.

The dominance of batch horizon, learning rate, and training iterations, with network width and batch size playing secondary roles, has direct practical consequences. Calibration workflows can therefore focus grid search on the three dominant hyper-parameters while fixing width to 5–15 neurons and keeping batch size low. The near-independence of pairwise interactions further supports sequential optimization rather than costly joint search. This empirical behavior is consistent with Theorem 1, since compact neural architectures restrict the effective flexibility of the neural correction term and help control the bounded hybrid deviation parameter *η* appearing in equation (14). Consequently, smaller networks simultaneously improve computational efficiency and strengthen preservation of mechanistic interpretability and parameter recoverability [49].

The reversal of NDE/HNDE performance ranking between the glycolysis (missing mechanism) and Lotka–Volterra (missing state) cases reveals that mechanistic constraints are beneficial only when the mechanistic form is correctly specified with respect to the dominant uncertainty. When the mechanistic scaffold correctly captures the dominant dynamics, HNDEs outperform pure NDEs in consistency. When state uncertainty dominates, however, the imposed structure becomes detrimental. Practitioners should therefore select the HNDE formulation based on the nature of the dominant uncertainty: additive neural corrections for unknown kinetics, and pure or lightly structured NDEs when missing states dominate.

HNDEs are natural building blocks for biological digital twins [50, 51]: computational replicas of biological systems that continuously assimilate data to refine predictions and guide interventions. The expressiveness–identifiability trade-off identified here has direct consequences for twin trustworthiness: a twin augmented with unconstrained neural dynamics may achieve high predictive fidelity while losing parameter interpretability, undermining its utility for mechanistic hypothesis testing and intervention design.

These results position hybrid neural dynamical systems as scientifically trustworthy tools for intelligent computing when identifiability-aware design principles are applied. Two directions follow directly. First, evaluating HNDEs on higher-dimensional biological systems and PDE-based models will test whether the expressiveness advantage grows with system complexity, as theory predicts. Second, integrating symbolic regression methods [42, 52] to extract interpretable closed-form expressions from trained neural components would close the loop between HNDE-based dynamic discovery and fully mechanistic modeling—completing the neural dynamics paradigm toward fully interpretable intelligent systems.

## Data availability

All code and data are publicly available at https://github.com/BenjaminWhipple/Hybrid-NODEs-Biology.git.

## Author contributions

**Benjamin Whipple:** Conceptualization, Methodology, Software, Formal analysis, Visualization, Writing—Original draft. **Esteban A. Hernandez-Vargas:** Conceptualization, Methodology, Formal analysis, Supervision, Funding acquisition, Writing—Review & editing.

